# Manganese influx and expression of ZIP8 is essential in primary myoblasts and contributes to activation of SOD2

**DOI:** 10.1101/494542

**Authors:** Shellaina J. V. Gordon, Daniel E. Fenker, Katherine E. Vest, Teresita Padilla-Benavides

**Affiliations:** Department of Biochemistry and Molecular Pharmacology, University of Massachusetts Medical School, 394 Plantation St., Worcester, MA, 01605, USA; S.J.V.G.; Department of Molecular Genetics, Biochemistry & Microbiology, University of Cincinnati School of Medicine, 231 Albert Sabin Way, Cincinnati, OH, 45267, USA

**Keywords:** myogenesis, manganese, ZIP8, ZIP14, Superoxide dismutase 2 (SOD2)

## Abstract

Trace elements such as copper (Cu), zinc (Zn), iron (Fe), and manganese (Mn) are enzyme cofactors and second messengers in cell signaling. Trace elements are emerging as key regulators of differentiation and development of mammalian tissues including blood, brain, and skeletal muscle. We previously reported an influx of Cu and dynamic expression of various metal transporters during differentiation of skeletal muscle cells. Here, we demonstrate that during differentiation of skeletal myoblasts an increase of additional trace elements such as Mn, Fe and Zn occurs. Interestingly the Mn increase is concomitant with increased Mn-dependent SOD2 levels. To better understand the Mn import pathway in skeletal muscle cells, we probed the functional relevance of the closely related proteins ZIP8 and ZIP14, which are implicated in Zn, Mn, and Fe transport. Partial depletion of ZIP8 severely impaired growth of myoblasts and led to cell death under differentiation conditions, indicating that ZIP8-mediated metal transport is essential in skeletal muscle cells. Moreover, knockdown of *Zip8* impaired activity of the Mn-dependent SOD2. Growth defects were partially rescued by Mn supplementation to the medium, suggesting additional functions for ZIP8 in the skeletal muscle lineage. Knockdown of *Zip14*, on the other hand, had only a mild effect on myotube size, consistent with a role for ZIP14 in muscle hypertrophy. This is the first report on the functional relevance of two members of the ZIP family of metal transporters in the skeletal muscle lineage, and further supports the paradigm that trace metal transporters are critical modulators of mammalian tissue development.

## INTRODUCTION

Transition metals such as copper (Cu), zinc (Zn), iron (Fe), and manganese (Mn), are essential for normal cellular function due to their roles as catalytic or structural cofactors and as signaling molecules. Transition metals also function as key regulators of mammalian development^1–8^. For example, Fe is a well-characterized regulator of erythropoiesis due to the high demand for heme biosynthesis in erythroid cells (reviewed by Ganz, *et al.*, 2012^9^). Recent studies have implicated other transition metals such as Cu in the differentiation of muscle, neuronal, and blood cells^1, 2, 10^. Like Cu and Fe, Mn acts as a catalytic or structural cofactor in enzymes involved in removal of oxygen radicals, gluconeogenesis, and protein glycosylation^11–13^. Excess Mn is toxic, and Mn accumulation leads to a neurodegenerative condition with symptoms similar to Parkinson’s disease and dystonia^14–16^. While few cases of dietary Mn deficiency have been reported, mutations in the Mn transporter, ZIP8, have been associated with skeletal and neuromuscular symptoms^17^. However, few studies have shown how Mn homeostasis and other transition metals impact skeletal muscle development.

Skeletal muscle largely consists of elongated, post-mitotic myofibers and a closely associated pool of stem cells, termed satellite cells^18–25^. Satellite cells are critical for embryonic and early postnatal muscle development, muscle regeneration, and may be important for maintenance of basal muscle function^26–28^. In response to injury or muscle damage, quiescent satellite cells are activated, proliferate as myoblasts, and differentiate into myocytes to regenerate skeletal muscle myofibers ^19–22, 24, 25, 29–38^. During this process, multiple metabolic changes occur to meet the fluctuating energetic demands. Quiescent satellite cells typically rely on fatty acid oxidation while proliferating myoblasts primarily utilize glycolysis for energy generation^37, 39^. During myoblast differentiation to myocytes, there is increased mitochondrial biogenesis and utilization of oxidative metabolism^39^. Given the importance of the mitochondrial Mn-superoxide dismutase (SOD2), Mn-influx and dynamic expression of Mn-transporting proteins are likely important factors in differentiation of skeletal muscle cells.

Several membrane metal transporters have been proposed to mobilize Mn in mammals^40–45^. Among these, two members of the family of Zrt- and Irt-like proteins (ZIP), ZIP8 (SLC39A8) and ZIP14 (SLC39A14), are considered the major Mn transporters in mammals^41, 46–48^. ZIP transporters have been mostly characterized for their function in importing Zn from the extracellular milieu to the cytosol^48–50^. They contain eight transmembrane helices (TM), form dimers, and have nanomolar to micromolar affinity for Zn^40, 46, 48, 51–53^. ZIP8 and ZIP14 have also been implicated in transport of Zn, Mn, Fe, Cu, and cadmium (Cd)^54^. These two proteins are the most evolutionarily closely related members of the LIV-1 subfamily of ZIPs, which contains the Zn-binding motif HEXXH in transmembrane domain 5 (TM5) (reviewed by Kambe *et al.*, 2015, and Jenkitkasemwong, *et al.*, 2015 ^48, 55^). Both ZIP8 and ZIP14 contain a conserved EEXXH motif in TM5; this single amino acid substitution may allow these proteins to transport other metals (reviewed by Jenkitkasemwong, *et al.*, 2015^55^). Thus, the specific metal binding residues in the TM domains of ZIP proteins may confer metal specificity. Despite the potential role for these ZIP transporters to mobilize a variety of ions, limited information is available regarding the participation of these metals and transporters in mammalian cell development, specifically in the skeletal muscle lineage. Importantly, our group demonstrated dynamic changes in the steady-state levels of mRNAs and proteins encoding the putative Mn transporters ZIP8 and ZIP14 during differentiation of immortalized muscle stem cell-derived C2C12 myoblasts^56^. Therefore, we hypothesized that a ZIP8- and/or ZIP14-mediated influx of intracellular Mn is necessary for myoblast differentiation.

We investigated the role of Mn in skeletal muscle differentiation using murine muscle stem cell-derived primary myoblasts. We detected increased cellular Mn in primary myoblasts starting at 24 h after inducing differentiation along with increased levels and activity of the manganese-dependent SOD2. We also detected increased expression of ZIP8 and ZIP14 during myoblast differentiation, suggesting that these two transport proteins are important mediators of metal transport in differentiating skeletal muscle cells. Partial loss of ZIP8 by shRNA-mediate knockdown led to cell death when myoblasts were induced to differentiate. ZIP8 knockdown also impaired proliferation of primary myoblasts, which supports an essential role for this transporter in the skeletal muscle lineage. Knockdown of *Zip14* partially impaired myotube formation but did not cause cell death indicating that ZIP14 contributes to, but is not essential for myogenesis. This study demonstrates that Mn, partially targeted to SOD2, is indispensable for the differentiation of the skeletal muscle lineage and that ZIP8 contributes to this process in primary cells. These results further support a critical role for transition metals in skeletal muscle differentiation, and this work lays the groundwork for future mechanistic studies to understand the tissue-specific transport functions of ZIP family proteins.

## EXPERIMENTAL METHODS

### Isolation, growth and differentiation of primary myoblasts

Mice were housed in the University of Massachusetts Medical School animal care facility as specified by Institutional Animal Care and Use Committee guidelines. Primary myoblasts derived from mouse satellite cells were isolated from hind limb muscles of wild type C57Bl/6 mice as previously described^30^. The myoblasts were grown in culture plates coated in 0.02% collagen (Advanced BioMatrix), in a 1:1 mix of DMEM and F-12 media (Gibco) supplemented with 20% fetal bovine serum, and 25 ng/ml recombinant basic FGF (Millipore). Myoblasts were induced to differentiate in DMEM containing 2% horse serum and 1% Insulin-Transferrin-Selenium-Sodium Pyruvate (Thermo Fisher Scientific). All cells were maintained in a humidified 5% CO_2_ incubator at 37 °C.

### Plasmids and lentivirus production

Mission plasmids (Sigma) encoding for three different shRNA against *Zip8* and *Zip14* (Table **S1**) were isolated by using the Pure yield plasmid midiprep system (Promega) following the manufacturer’s instructions. shRNA (15 μg) and the packing vectors pLP1 (15 μg), pLP2 (6 μg), pSVGV (3 μg) were transfected using lipofectamine 2000 (Thermo Fisher) into HEK293T cells for lentiviral production. After 24 and 48 h the supernatants containing viral particles were collected and filtered using a 0.22 μm syringe filter (Millipore). Primary myoblasts were transduced with lentivirus in the presence of 8 mg/ml polybrene and selected with 1.5 μg/ml puromycin (Invitrogen).

### Whole cell Mn and Zn content analysis

Three independent biological replicates of proliferating and differentiating (24, 48, and 72 h) primary myoblasts were rinsed three times with PBS without Ca^+2^ and Mg^+2^ (Gibco). Cells were resuspended in 100 μl PBS and lysed by sonication using a Diagenode Bioruptor UCD-200. Total protein content was quantified using Bradford assay^57^. The samples were resuspended in concentrated HNO_3_ (trace metal grade) and analyzed by inductively coupled plasma-optical emissions spectroscopy (ICP-OES) as previously described^58^. Concentrations of Mn, Zn, Fe, and Ca measured via ICP-OES were normalized to the initial mass of protein in each sample.

### Western blot analyses

Protein samples from proliferating and differentiating primary myoblasts (control and *Zip8*- and *Zip14*-knockdown) were solubilized with RIPA buffer (10 mM piperazine-N,N-bis(2-ethanesulfonic acid), pH 7.4, 150 mM NaCl, 2 mM ethylenediamine-tetraacetic acid (EDTA), 1% Triton X-100, 0.5% sodium deoxycholate and 10% glycerol) supplemented with Complete protease inhibitor cocktail (Roche). Protein content was quantified by Bradford^57^. Samples (30 μg) were separated by SDS-PAGE and electrotransferred to PVDF membranes (Millipore). The proteins of interest were detected using the primary antibodies rabbit anti-ZIP8 (A10395), anti-ZIP14 (A1043), anti-SOD1 (A0274), anti-SOD2 (A1340) anti-Caspase 3 (A2156) from Abclonal. Mouse anti-actin (sc-8432) from Santa Cruz Biotechnologies was used as a loading control. Secondary antibodies used were horseradish peroxidase-conjugated anti-mouse and anti-rabbit (Thermo Fisher). Chemiluminescent detection was performed using ECL PLUS (GE Healthcare).

### SOD Activity Gels

Protein samples from proliferating and differentiating primary myoblasts (control and *Zip8*- and *Zip14*-knockdown) were solubilized with RIPA buffer plus protease inhibitor cocktail and quantified by Bradford assay^57^. Fifty micrograms of each sample were separated using native PAGE gels. Gels were incubated for 30 min in 2.5 mM nitro blue tetrazolium (Sigma), followed by a 20-min incubation with 30 mM potassium phosphate, 30 mM TEMED, 30 mM riboflavin, pH 7.8^59^. Superoxide dismutase activity was visualized by illuminating gels with white light for 10 min on a transilluminator. Areas of activity were visible as white bands against a dark background. SOD2 activity was detected as a cyanide-insensitive 26 kDa band, while SOD1 activity was detected as a 17 kDa band and a series of higher molecular weight oligomers (Supp. Fig. **4**).

### Immunocytochemistry

Proliferating and differentiating primary myoblasts (control and *Zip8*- and *Zip14*-knockdown) were fixed overnight in 10% formalin-PBS at 4 °C. Samples were washed with PBS and permeabilized for 10 min in PBS containing 0.2% Triton X-100. Immunocytochemistry was performed using universal ABC kit (Vector Labs) following manufacturer’s instructions. Hybridoma supernatants from the Developmental Studies Hybridoma Bank (University of Iowa) were used against myogenin (F5D, deposited by W.E. Wright), and Pax7 (deposited by A. Kawakami).

### Statistical analysis

All statistical analyses were performed using Kaleidagraph (Version 4.1). Statistical significance was determined using one-way analysis of variance (ANOVA), followed by Bonferroni multiple comparison tests. Experiments where P<0.05 were considered statistically significant.

## RESULTS

### Manganese and Mn-dependent SOD2 accumulate in differentiating myoblasts

Previous reports from our group have shown that the concentrations of intracellular Zn and Cu increase during myogenic differentiation^2, 56^. To better understand how additional metals fluctuate during myogenic differentiation, we utilized ICP-OES to measure total metals in proliferating and differentiating primary myoblasts. We detected an increase cellular Mn starting 24 h after inducing differentiation (Figure **1A**). We also detected a slight non-significant increase in cellular Fe after three days of differentiation (Figure **1B**). Consistent with our recently reported study of zinc in immortalized C2C12 myoblasts^56^, we detected a slight increase in intracellular Zn 24 h after inducing differentiation in primary myoblasts (Figure **1C**). In all cases, intracellular Ca measurements were used as a positive control for differentiated muscle cells (Figure **1D**).

**Figure 1.**
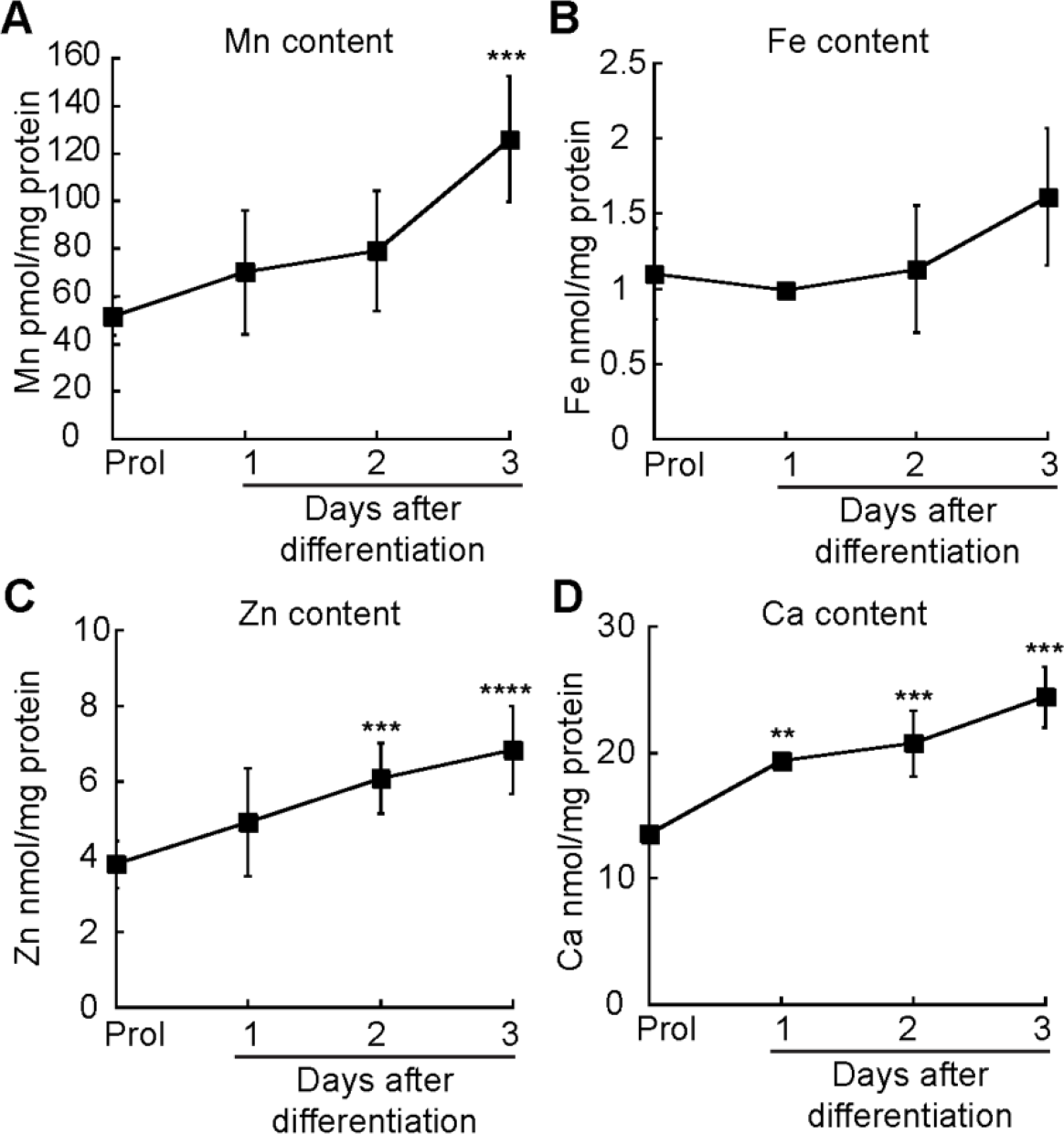
Metal intake in differentiating primary myoblasts. Whole cell metal content analyses of proliferating and differentiating wild type myoblasts. (**A**) Manganese. (**B**) Iron. **(C)** Zinc. (**D**) Calcium. All data were obtained using ICP-OES and normalized to total protein. Statistical analyses showed significant differences in metal accumulation in differentiating myoblasts compared to proliferating cells. Data is the mean ± standard error for three independent biological replicates. ****P<0.001, ***P < 0.005, **P < 0.01

During differentiation, myoblasts switch from primarily glycolytic to more oxidative metabolism^39, 60–62^. This is associated with a concomitant increase in mitochondrial biogenesis and increased reactive oxygen species^39, 60–62^. Therefore, the mitochondrial Mn-dependent SOD2 is a likely target of the increased influx of intracellular manganese during myogenic differentiation. Immunoblot analysis and in-gel activity staining revealed an increased level of SOD2 protein and activity during differentiation of primary myoblasts when compared to proliferating cells (Figure **2**). Importantly, SOD2 was distinguished from SOD1, the Cu/Zn superoxide dismutase by potassium cyanide (KCN) inhibition^63^ (Supp. Figure. **1**). These data suggest that SOD2 is a relevant target for Mn during myogenic differentiation.

**Figure 2.**
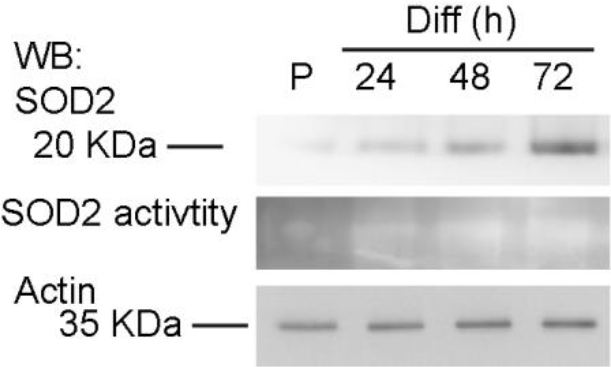
SOD2 expression and activity increases as a consequence of myogenic differentiation. Representative immunoblots and activity gels analyses for SOD2 in proliferating and differentiating myoblasts expressing, Actin and Coomasie-stained membranes were used as loading controls (Supp. Figure **9**).

### ZIP8 and ZIP14 transporters are induced during the differentiation of primary myoblasts derived from mouse satellite cells

To better understand how Mn is transported into primary myoblasts during differentiation, we looked to the ZIP family of metal transporters. In particular, ZIP8 and ZIP14 are two homologous Zn, Mn and Fe transporters, which share 46.6% of identity and 62.0% similarity (Supp. Fig. **2**). Previous investigations from our laboratory showed that in immortalized C2C12 myoblasts, expression of ZIP8 was induced during myogenesis, while no change was detected in the levels of ZIP14^56^. However, chromosomal alterations found in immortalized cell lines such as C2C12 cells can confound data. Therefore, we sought to study expression of ZIP8 and ZIP14 in primary myoblasts derived from mouse hind limb muscle stem cells. To determine whether the increase in ZIP8 was recapitulated in primary myoblasts, we performed immunoblots using antibodies to ZIP8 and ZIP14 in proliferating cells and cells differentiated for 24, 48, and 72 h. As shown in Figure **4A**, levels of ZIP8 increased by approximately 50% starting 24 h after induction of differentiation and more than doubled after 72 h of differentiation. This result is similar to the increase detected in differentiating C2C12 myoblasts^56^. Levels of ZIP14 showed a small increase 48 and 72 hours after differentiation (Fig. **4B**). These results indicate that, unlike ZIP14 in C2C12 myoblast differentiation, levels of ZIP14 increase during later stages of primary myoblast differentiation.

### *Zip8* and *Zip14* knockdowns have differential effects on the differentiation of primary myoblasts

Given the increased expression of ZIP8 and ZIP14 observed during primary myoblast differentiation (Figure **4**) and the roles of these proteins in Mn transport^41, 64, 65^, we hypothesized that ZIP8- and/or ZIP14-dependent Mn transport may contribute to the maturation of the skeletal muscle lineage. To determine the role of ZIP8 and ZIP14 in myoblast differentiation, we used viral vectors encoding small hairpin RNAs (shRNA) to knockdown these two transporters in primary myoblasts. Three lentiviral constructs (Supp. Table **1**) that encode shRNAs against *Zip8* and *Zip14* mRNA were used to knockdown the endogenous transporters in proliferating and differentiating primary myoblasts. Myoblasts transduced with a lentivirus-encoded non-specific scramble shRNA were used as negative controls. The virus-infected cells were selected with puromycin and the levels of *Zip8* and *Zip14* were examined by Western blot analysis (Figure **3**, Supp. Fig. **4**). Cells containing *Zip8* shRNA failed to increase expression of ZIP8 protein upon incubation in differentiation medium (Figure **3A**, Supp. Figure **3A**). Cells containing *Zip14* shRNA showed increased levels of ZIP14 during differentiation but to a much lesser degree and at a much later time point than wild type cells (New Figure **3B**, Supp. Figure **3B**).

**Figure 3.**
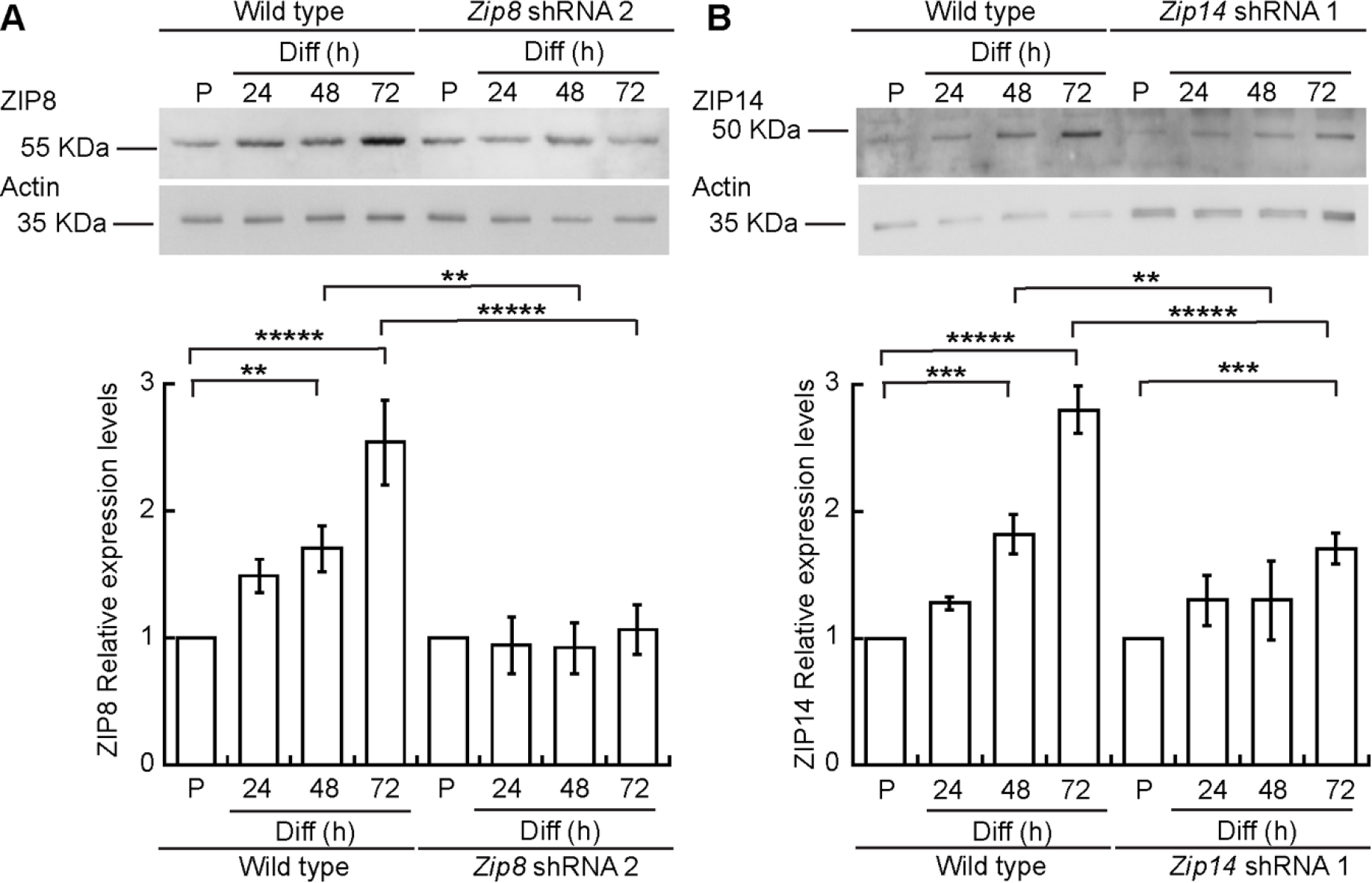
ZIP8 and ZIP14 expression in proliferating and differentiating wild type and shRNA knockdown primary myoblasts derived from mouse satellite cells. **(A)** Representative immunoblot (top) and quantification (bottom) of ZIP8 levels in proliferating myoblasts and differentiated cells at 24, 48, and 72 h post differentiation in wild type (left side panel) and *Zip8* shRNA expressing cells (right side panel). **(B)** Representative immunoblot of ZIP14 levels in proliferating and differentiating wild type (left) and *Zip14* shRNA myoblasts. For all samples, shown is mean ± standard error of three independent biological replicates. Immunoblots against actin or Coomasie-stained membranes (Supp. Fig. **8**) were used as loading controls. Samples were compared to the corresponding wild type or mutant proliferation time point, and the; mutants were compared also to the equivalent time point in wild type cells. *****P<0.0001 ****P<0.001, ***P < 0.005, **P < 0.01

To determine whether the partial loss of ZIP8 or ZIP14 impaired myogenesis, *Zip8* and *Zip14* shRNA-treated primary myoblasts were grown to confluence and induced to differentiate by serum deprivation^66^. Immunocytochemical staining and immunoblot analyses for the lineage specific differentiation marker myogenin was used as an indicator of proper myogenesis in samples taken during the proliferation stage and at 24, 48 and 72 h post-differentiation (Figure **4;** Supp. Figure **4** and **5**). *Zip8* knockdown cells did form elongated myocytes 24 h after inducing differentiation, but also detached from the plates and failed to form myotubes at later time points (Figure **4A**; Supp, Figure **4A** and **5**). The *Zip8* knockdown cells had a decreased frequency of myogenin positive nuclei and showed a significantly lower fusion index than wild type cells at similar time points (Figure **4A, B**, Supp. Figure **4, 5**). Cells containing *Zip14* shRNA formed myotubes but also showed a lower fusion index than control cells at all time points after inducing differentiation (Figure **4A, B**, Supp. Fig. **4, 5**). Importantly, these results suggest that ZIP14 contributes to robust myotube formation, but ZIP8 is essential for differentiation.

**Figure 4.**
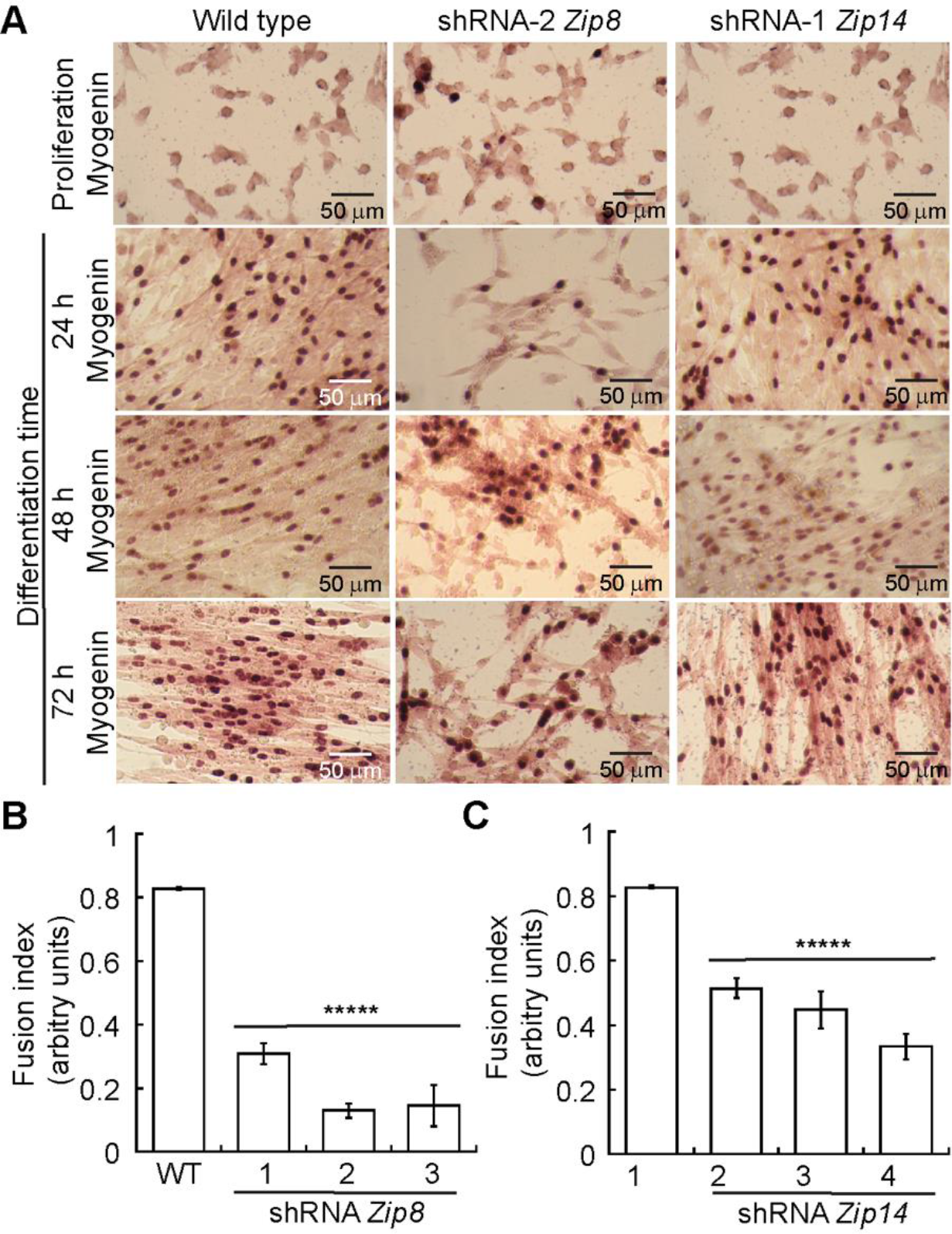
Knockdown of *Zip8* impairs differentiation of primary myoblasts. **(A)** Representative light micrographs of wild type, *Zip8* and *Zip14* knockdown myoblasts grown in proliferating conditions or at 24, 48 and 72 h after inducing differentiation. Cells were immunostained with an anti-Myogenin antibody. Calculated fusion index for *Zip8*- (**B**) and *Zip14*-shRNA treated myoblasts. (**C**) Data represent mean ± SE for three independent experiments. *****P<0.0001

The detachment detected in *Zip8* knockdown cells after inducing differentiation suggests that partial loss of ZIP8 leads to cell death. A major cell death pathway is caspase-3-dependent apoptosis^67^. Myoblast differentiation requires the disassembly and reorganization of actin fibers and the activation of myosin light chain kinase, events that also occur during apoptosis^68–70^. Previous studies have shown that myogenic differentiation and apoptotic processes undergo similar mechanisms of cleaving and activation of caspase 3^71–74^. We analyzed by immunoblot the cleavage of Caspase 3 in proliferating and differentiating primary myoblasts. We found the expected low levels of activated caspase 3 in wild type proliferating and differentiating myoblasts (Supp. Figure **5B**). Furthermore, we observed an increase in the active form of caspase 3 48 h after inducing myogenesis in cells transduced with the three different *Zip8* shRNA (Supp. Figure **5B**). No changes in the levels of activated caspase 3 were detected in the *Zip14*-treated shRNA myoblasts, which is consistent with the fact that these cells did not detach upon inducing differentiation (Supp. Figure **5B**).

### *Zip8* knockdown cells fail to expand intracellular Mn and Mn-dependent SOD2 after being induced to differentiate

Both ZIP8 and ZIP14 are have been implicated in transport of Mn, Zn and Fe (reviewed by Jenikitkasemwong, *et al.*, 2015^55^), which are critical for cell growth and survival^41^. To understand how partial loss of ZIP8 and ZIP14 affect accumulation of Mn, Fe, and Zn, we used ICP-OES to measure total metal quotas in differentiating *Zip8* and *Zip14* knockdown cells (Figure **5**). Similar to wild type cells (Figure **1A**), myoblasts transfected with scrambled (scr) shRNA showed a significant increase in intracellular Mn starting at 24 h after inducing myogenic differentiation (Figure **6A**). Cells transduced with *Zip8* shRNA failed to accumulate Mn while those transduced with *Zip14* shRNA showed no defect in Mn accumulation (Figure **5A**; Supp. Figure **6**). Measurement of total levels of Fe (Figure **5B**; **Supp.** Figure **6**), Zn (Figure **5C;** Supp. Figure **6**) revealed similar results, where wild type and *Zip14* knockdown cells showed accumulation at 24 h after inducing differentiation, but *Zip8* knockdown cells did not. Measurement of total Ca was used as a positive control for myogenic differentiation (Figure **5D**; Supp. Figure **7**). Total calcium levels increased cells transfected with scrambled shRNA and *Zip14* shRNA at 24 h after inducing differentiation (Figure **5C**; Supp. Figure **7**). However, *Zip8* knockdown cells failed to accumulate Ca after being induced to differentiate (Figure **5C;** Supp. Figure **7**). These results are consistent with the severe differentiation defect observed in *Zip8* knockdown cells (Figure **4**; Supp. Figure **5**).

**Figure 5.**
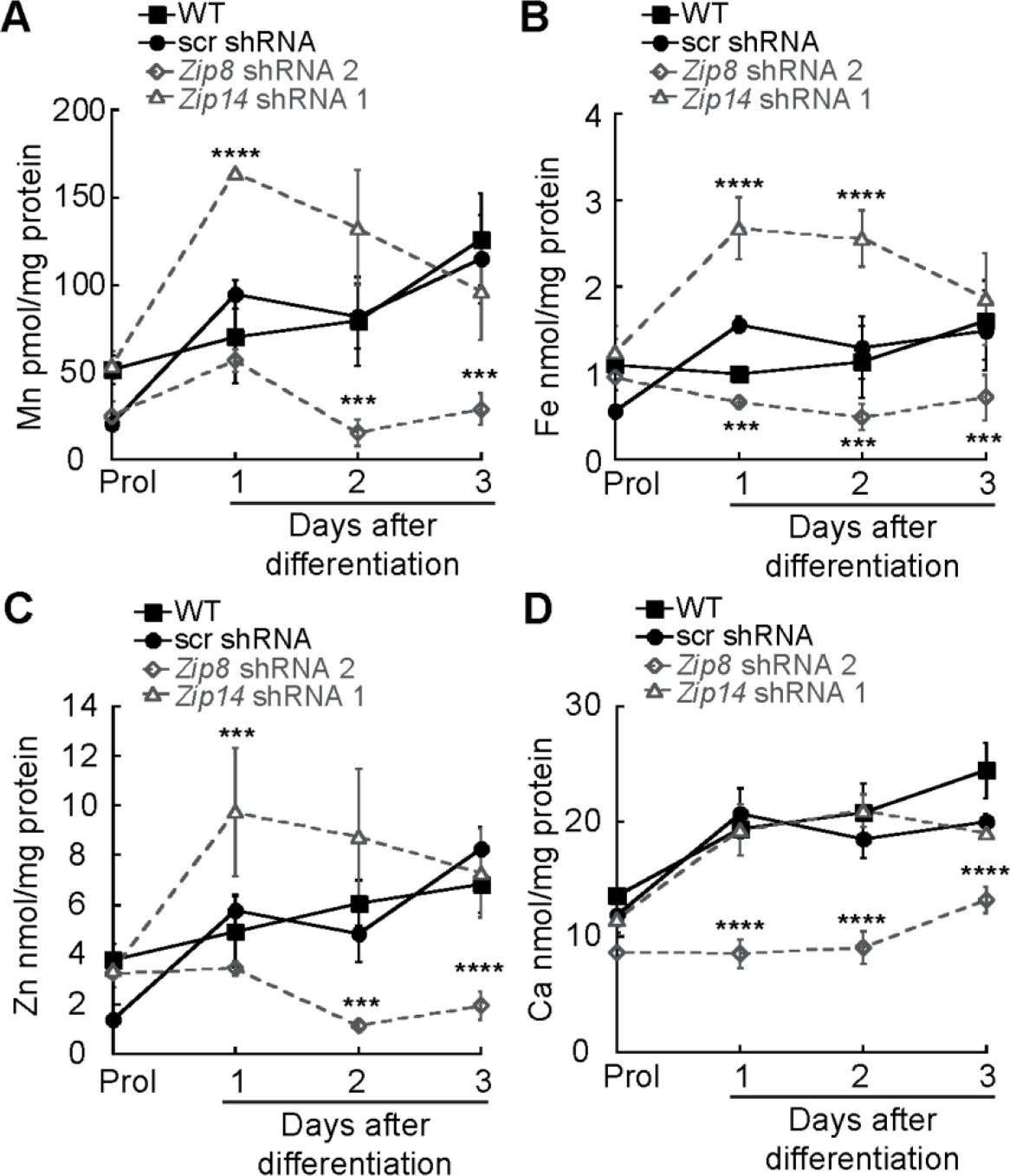
Knockdown of *Zip8*, but not *Zip14* impairs metal intake in differentiating primary myoblasts. Whole cell metal content analyses of proliferating and differentiating wild type or myoblasts transduced with scrambled (scr), *Zip8* or *Zip14* shRNA. Statistical analyses showed significant differences in metal accumulation in differentiating myoblasts (**A**) Manganese. (**B**) Iron. **(C)** Zinc. (**D**) Calcium. All data were obtained using ICP-OES and normalized to total protein. Shown is mean ± standard error for three independent biological replicates. ****P<0.001, ***P < 0.005

**Figure 6.**
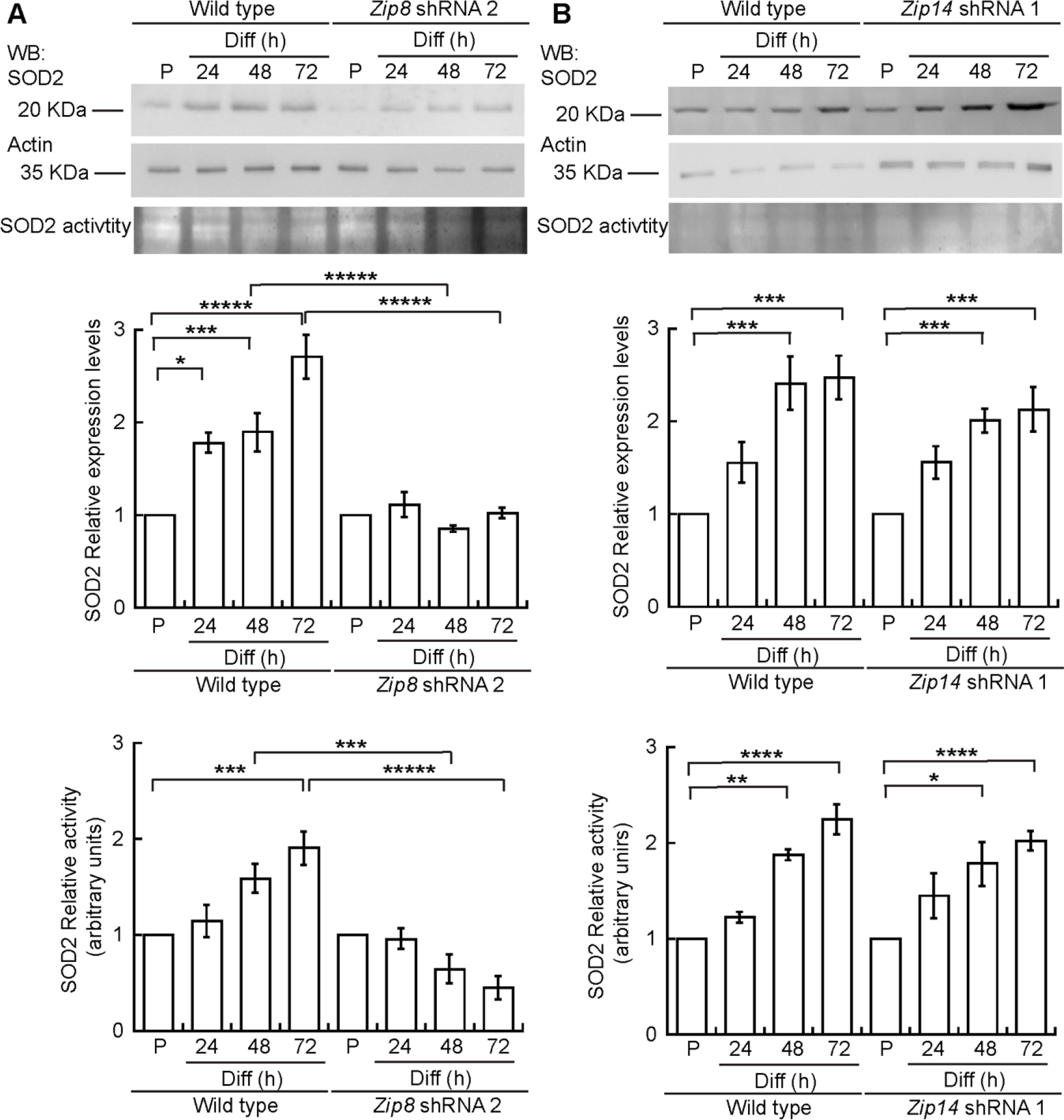
SOD2 expression and activity is decreased in *Zip8* but not *Zip14* knockdown primary myoblasts. **(A)** Representative Western blots and activity gels (top panels) and quantification of SOD2 levels and activity (bottom panels) in wild type and *Zip8* shRNA cells. **(B)** Representative Western blots and activity gels (top panels) and quantification of SOD2 levels and activity (bottom panels) in wild type and *Zip14* shRNA cells. For all samples, blots against actin and Coomassie-stained membranes were used as loading controls (Supp. Fig. **8**). Shown is mean ± standard error for three biological replicates. For wild type differentiating myoblasts, statistical analyses showed significant differences when compared to proliferating cells. Bonferroni statistical analyses for *Zip8*-knock down cells showed significant differences when compared to control cells at the corresponding time points, and also when compared to the mutant’s proliferation point. Similar to control cells, *Zip14*-knockdown cells showed significant differences when compared to proliferating *Zip14*-mutant myoblasts; *****P<0.0001 ****P<0.001, ***P < 0.005, **P < 0.01, *P ≤ 0.05

ZIP8 has been previously shown to transport Mn^44^, which is a cofactor for the mitochondrial SOD^75^. SOD2 has a well-characterized function in maintaining redox balance and mitochondrial function in skeletal muscle as well as in promoting proliferation and differentiation of myoblasts^76–78^. To determine whether differentiation defects in *Zip8* knockdown myoblasts are associated with a loss of SOD2 activity, we performed immunoblot and *in-gel* activity assays of SOD2. As shown in figure 2, an increased steady-state level and activity of SOD2 was detected in wild type myoblasts induced to differentiate. Therefore we asked whether the partial knockdown of ZIP8 or ZIP14 would impair the expression or activity of SOD2. In the *Zip8* knockdown myoblasts, the levels of SOD2 protein and activity failed to increase as a consequence of the myogenic differentiation program (Figure **6A**; Supp. Fig. **8A, C**). SOD2 protein levels and activity in proliferating and differentiating *Zip14* knockdown myoblasts were similar to wild type and scramble control (Figure **6B**, Supp. Fig. **8B, D**). Protein levels and activity of SOD2 was normalized to total protein (Supp. Fig. **9**). These results could support a role for ZIP8, but not ZIP14, in SOD2 activation in differentiating myoblasts. However, impaired SOD2 activity could also be a result of impaired differentitation in *Zip8* knockdown cells.

### Knockdown of *Zip8*impairs proliferation of primary myoblasts

Considering that knockdown of *Zip8* in primary myoblasts led to cell death after inducing differentiation and that Mn, Fe, and Zn are important for cellular growth and survival, we sought to understand how myoblasts proliferation is affected. We used cell counting to as a proxy measurement for cell proliferation rate in *Zip8* knockdown cells. While numbers of wild type and myoblasts transduced with scrambled shRNA increased 50 fold over three days (Figure **7**, Supp. Figure **10**), cells containing *Zip8* shRNA increased in number only 10 fold over three days (Figure **7**; Supp. Figure **10**). Expression of PAX7 was normal in proliferating *Zip8* knockdown cells, indicating that growth defects are not related to loss of this myoblast-specific transcription factor (Figure 7 E,F). To determine whether impaired Mn import is responsible for the cellular growth defect detected in *Zip8* knockdown cells, we supplemented the growth medium with increasing concentrations of Mn (Supp. Figure **11**). Addition of 300 μM Mn to the growth medium led to a mild decrease in proliferation of shRNA scrambled and wild type cells (Supp. Figure **11A**) but partially rescued the growth defects detected in *Zip8* knockdown cells (Fig. **8**; Supp. Figure **11B**). Cell counting assay revealed that control and Mn-treated cells expressing *Zip14* shRNA-1 proliferated similarly to wild type and shRNA scrambled cells (Supp. Figure **11C**). These results indicate that partial loss of ZIP8 impacts myoblast proliferation and can be moderately rescued by supplementation with exogenous Mn, further supporting the conclusion that ZIP8 is essential in this cell type.

**Figure 7.**
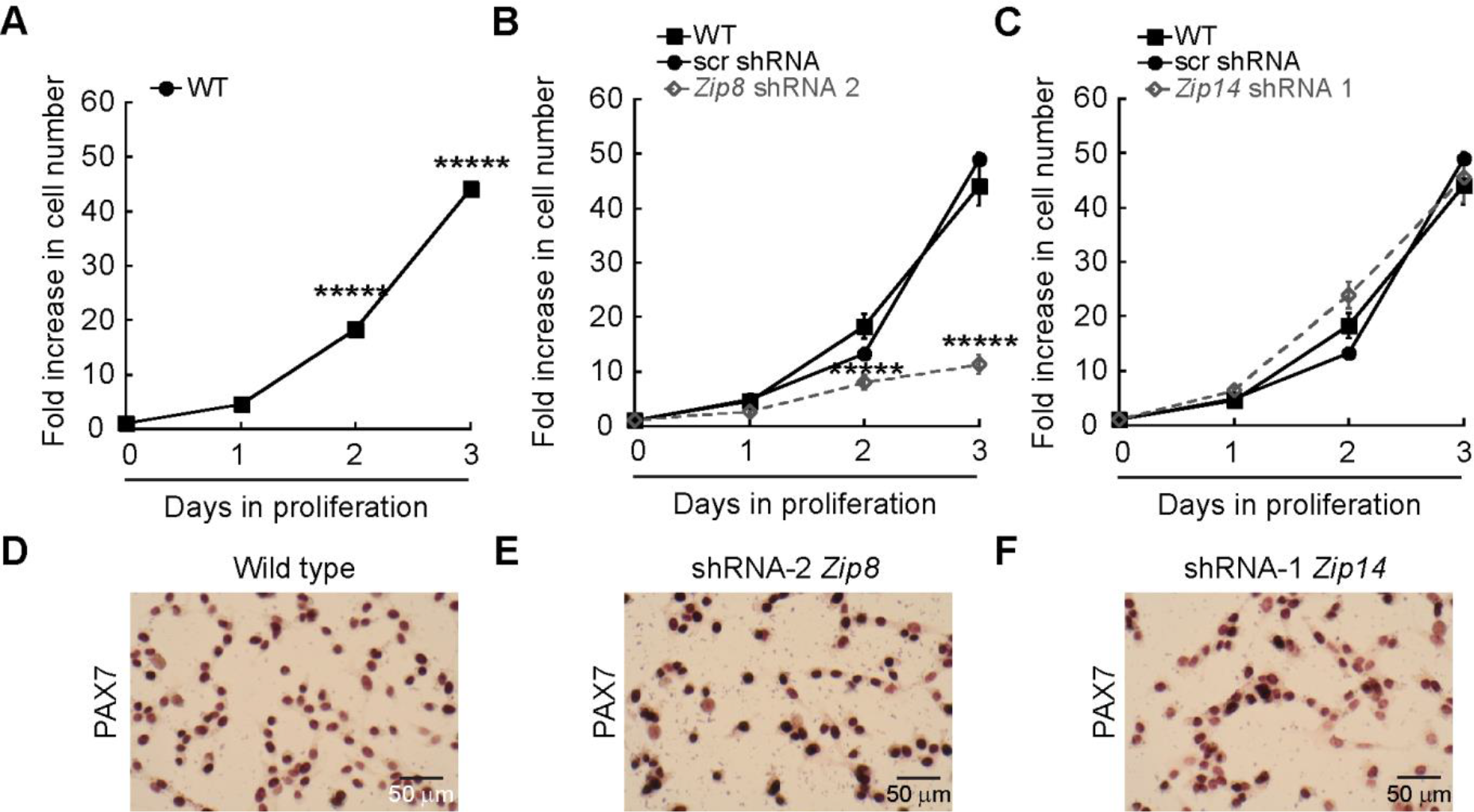
Partial depletion of *Zip8*, but not *Zip14*, impairs growth of primary myoblasts. **(A-C)** Cell counting assay of proliferating wild type myoblasts, myoblasts transduced with scrambled shRNA (shRNA scr, B,C), *Zip8* (**B**) or *Zip14* shRNAs (**C**). Data in A-C are mean ± standard error for three independent experiments. *****P<0.0001. **(D-F)** Representative light micrographs of proliferating wild type myoblasts (**D**), *Zip8* (**E**) or *Zip14* (**F**) knockdown myoblasts immunostained for Pax7.

**Figure 8.**
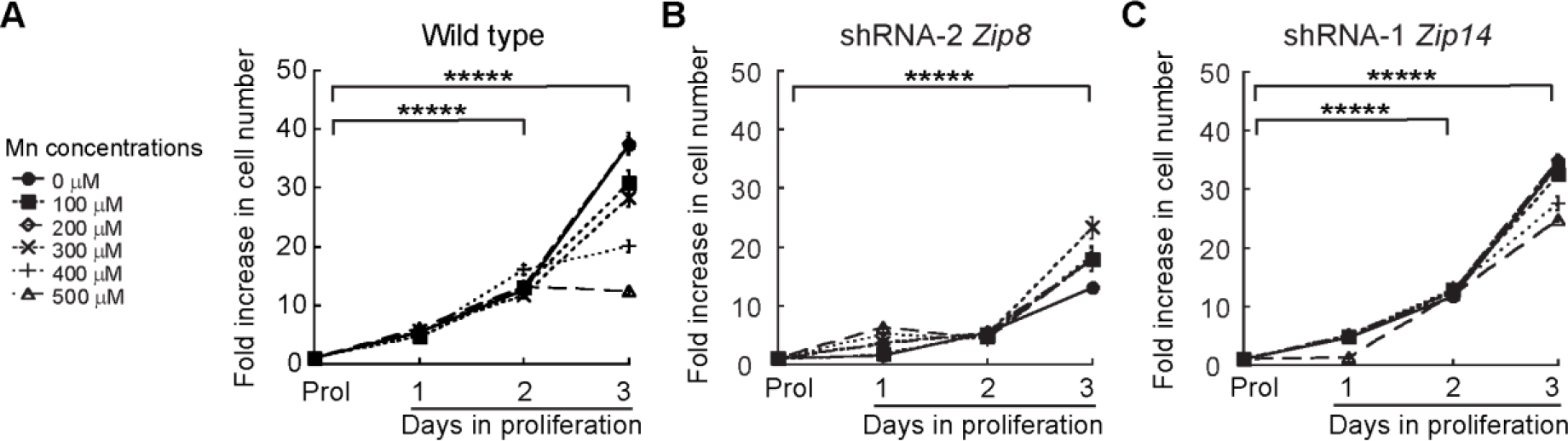
Mn partially rescues the growth defect observed in knockdown *Zip8* primary myoblasts. **(A)** Cell counting assay of proliferating wild type myoblasts grown in increasing concentrations of Mn. **(B)** Cell counting assay of *Zip8* knockdown myoblasts grown in increasing concentrations of Mn. **(C)** Cell counting assay of *Zip14* knockdown myoblasts grown in increasing concentrations of Mn. For all experiments, data represent mean ± standard error for three independent experiments. *****P<0.0001

## DISCUSSION

Myogenesis encompasses metabolic and morphological changes that largely depend on the bioavailability of transition metals to enable energy production and redox homeostasis^79, 80^. Here, we show that the concentration of Mn increases within 24 h of inducing differentiation in primary myoblasts. This influx of Mn is associated with increased expression and activity of SOD2, which is consistent with increased mitochondrial biogenesis in differentiating myoblasts^39, 60, 61^. Given that the plasma membrane is a critical barrier for metal influx into cells, we sought to understand the functional relevance of metal transporters in primary myogenesis. We focused on ZIP8 and ZIP14 due to their function in Mn, Fe, and Zn transport^40, 41, 44, 64, 65^. We detected a one to two-fold increase in expression of ZIP8 as early as 24 h of initiation of differentiation of primary myoblasts, which was consistent with our previous findings from C2C12 cells^56^. We also detected a small and sustained increase in the expression of ZIP14 after 48 to 72 h of differentiation. The detected increase in ZIP14 differs from our previous report showing that ZIP14 protein levels do not change during differentiation of C2C12 myoblasts^56^, which can be explained by the intrinsic limitations of immortalized cell lines.

To determine the functional significance of ZIP8 expression, we used viral vectors encoding shRNAs to knockdown *Zip8* in primary myoblasts. Attempts to differentiate *Zip8* knockdown cells were unsuccessful as cells failed to form myotubes and showed no increase in the differentiation marker myogenin. We observed a failure in the differentiation-dependent increase in Mn levels and Mn-dependent SOD2 expression and activation in the *Zip8-shRNA* transduced myoblasts. However, this failure to import Mn could be an indirect result of failed differentiation or cell death. Indeed, we detected increased levels of cleaved caspase-3, a marker of apoptosis, in *Zip8* knockdown cells after inducing myogenesis. These results suggest that ZIP8 may be essential for growth and survival in primary myoblasts, which is supported by the fact we detected proliferation defects in *Zip8* knockdown cells. The proliferation defect detected after partial depletion of ZIP8 was only partially reversed by addition of exogenous Mn. The lack of complete rescue of the *Zip8* knockdown growth defect by Mn suggests that ZIP8 is important for transport of additional substrates in skeletal muscle cells. Previous studies have demonstrated that ZIP8 expression in skeletal muscle is low relative to other tissues, and that depletion of ZIP8 does not cause overt muscle damage^81, 82^. However, no previously published data demonstrate the functional importance of ZIP8 in primary myoblasts. Interestingly, several reports have demonstrated a connection between ZIP8 and proliferation and survival of cancer cells^83, 84^. Further studies are needed to determine whether ZIP8-mediated metal transport is a general requirement for proliferation and survival in mammalian cells.

Partial depletion of ZIP8 resulted in decreased levels and activity of SOD2 in proliferating cells. A previous study utilizing *Sod2*^+/−^ myoblasts showed that partial loss of SOD2 function impaired both proliferation and differentiation, but not to the degree detected in this study^78^. The discrepancy between *Sod2*^+/−^ and *Zip8* knockdown myoblasts supports a model where ZIP8 performs other important functions in addition to import of Mn for SOD2. This model is reinforced by our observation that supplementation of exogenous Mn in the medium only partially reversed the growth defect in proliferating *Zip8* knockdown myoblasts. For instance, ZIP8 expression has been correlated with activation of the metal-sensing transcription factor (MTF1), which has important functions in myoblasts such as inducing the transcription of genes important in skeletal muscle including matrix metalloproteases^85^. Additional transport substrates of ZIP8 in myoblasts remain to be elucidated. Although we detected decreased levels of Mn, Zn, Fe, in *Zip8* knockdown cells, we cannot overrule the possibility that this decrease is an indirect effect of the severe growth defect in *Zip8* knockdown cells.

To further characterize metal transport in differentiating myoblasts, we also sought to determine the functional relevance of the closely related transporter, ZIP14. Cells partially depleted of ZIP14 showed no proliferation defects and formed myotubes in differentiation medium. At 24 and 48 h after differentiation, the *Zip14* knockdown myotubes resembled those in formed by control cells. However, by 72 h after inducing differentiation, it was apparent that *Zip14* formed fewer and smaller myotubes as detected by analysis of fusion index. The presence of normal differentiation but smaller myotubes suggests that ZIP14-mediated metal transport contributes to myotube hypertrophy rather than myoblast differentiation. Given the function of ZIP14 in Zn transport, our results are consistent with a previous study showing that increased intracellular Zn availability, initiated either by metallothionein knockout or treatment with zinc-pyrithione, promotes muscle hypertrophy^86^. However, a recent report has demonstrated that elevated ZIP14-mediated Zn transport impairs myoblast differentiation and promotes cachexia in mouse cancer models^87^. It is likely a dual scenario where the dosage of ZIP14 is important for normal muscle function and that too little or too much is deleterious. We did not detect changes in the accumulation Mn, Fe, or Zn in *Zip14* knockdown cells. Instead, metals levels were highly variable depending on which shRNA was utilized and some samples appeared to accumulate more Mn, Zn and Fe. Future studies are needed to further elucidate the specific transport substrates of ZIP14 in myoblasts.

Mutations in human *ZIP14* have been linked to symptoms of the early onset of Parkinsonism and dystonia^16^. However, this phenotype has been proposed to be related to Mn accumulation in the nervous system, rather than direct effects in skeletal muscle^16^. Expression of ZIP14 is induced during inflammation and skeletal muscle is one of several tissues where this transporter is overexpressed as a consequence of lipopolysaccharide insult^88^. The early stages of muscle regeneration involve an M1-macrophage mediated pro-inflammatory response (reviewed by Juban *et al.*, 2017^89^), so it is plausible that ZIP14 expression is important in early stages of muscle stem cell activation. Data from an RNA-seq study comparing quiescent and activated satellite cells showed that steady-state levels of *Zip14* mRNA increase two-fold in newly activated satellite cells relative to quiescent satellite cells^90^. Interestingly, levels of *Zip8* mRNA are also upregulated during early activation^90^, suggesting that these two transporters work cooperatively to mediate metal transport in activated muscle stem cells. Indeed, the subtle phenotype detected in *Zip14* knockdown cells would support a model in which metal transport via ZIP8 can partially compensate for the loss of ZIP14. However the mechanistic details and implications potential genetic interactions between *Zip8* and *Zip14* remain to be elucidated.

## CONCLUSIONS

Our work is the first study demonstrating manganese influx and SOD2 metallation in the context of skeletal muscle differentiation. We found that ZIP8 is essential in primary myoblasts derived from mouse satellite cells, and that ZIP14 plays a non-essential role in myotube hypertrophy. Future studies are needed to determine the mechanistic details of interactions between ZIP8 and ZIP14 and the specific metallo-substrates of each transporter in myoblasts.

## Supporting information

## ACKNOWLEDGMENTS

This research was supported by the Faculty Diversity Scholars Award from the University of Massachusetts Medical School, to T.P.-B. S.J.V.G. was supported by the University of Massachusetts Medical School funding for the Summer Undergraduate Research Experience program. The authors thank Dr. Paul Cobine (Auburn University) for assistance with ICP-OES experiments and Ms. Daniella Cangussu for her technical assistance.

## AUTHOR CONTRIBUTIONS

T. P.-B. and K. E. V: Conception and design; T.P.-B, K.E.V, S.J.V.G and D. E. F.: Acquisition of data, Analysis and interpretation of data, Drafting and revising the article.

## CONFLICTS OF INTEREST

The authors declare no conflict of interest. The founding sponsors had no role in the design of the study; in the collection, analyses, or interpretation of data; in the writing of the manuscript, and in the decision to publish the results.

