## Supplementary material for "Manganese influx and expression of ZIP8 is essential in primary myoblasts and contributes to activation of SOD2"

#### SUPPLEMENTARY INFORMATION

#### SUPPLEMENTARY FIGURES

##### Supplementary figure 1

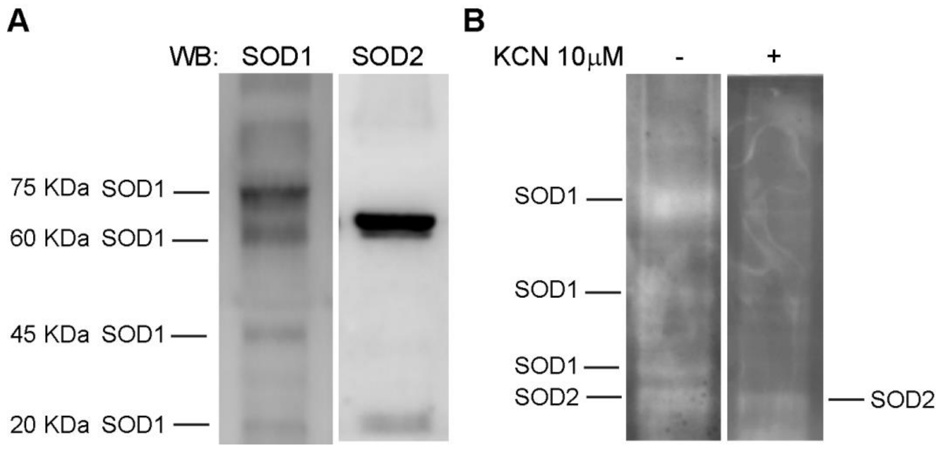


**Supplementary Figure 1. In-gel assays for SOD1 and SOD2 activity.** Representative figure showing how SOD1 and SOD2 activities were differentiated on native gels. **(A)** Immunoblot on a native gel loaded with cell lysates from proliferating wild type myoblasts and probed with antibodies against SOD1 and SOD2. **(B).** In-gel activity assay showing gel probed with nitro-blue tetrazolium on gels with and without cyanide treatment to strip the Cu cofactor from SOD1.

#### Supplementary figure 2

```
ZIP8          1 ---------------------MAP--GRAVAGLLLLAATSLGHPSEGPEL   27
                                     .||  ..:.|||..|:|||
ZIP14         1 MKRLHPALPSCLLLVLFGIWRTAPQTHASSAGLPPLSATS----------   40

ZIP8         28 AFSEDVLSVFGANRSLSAAQLGRLLERLGAASQQGALDL---GQLHFNQC   74
                 |.||::..:|.|.||:..||..||:.|.....:..:..   |..:.:.|
ZIP14        41 -FLEDLMDRYGKNDSLTLTQLKSLLDHLHVGVGRDNVSQPKEGPRNLSTC   89

ZIP8         75 LSAEDIFSLHGFSNVTQITSSNFSAICPAILQQLNFHPC--------EDL  116
                .|:.|:|:.|..|..:||.:|.|...||.|||||:...|        |:.
ZIP14        90 FSSGDLFAAHNLSERSQIGASEFQEFCPTILQQLDSQACTSENQKSEENE  139

ZIP8        117 RKHNAKPSLSEVWGYGFLSVTIINLASLLGLILTPLIKKSYFPKILTYFV  166
                :....|||..||||||||.||:|:|.||:|..:.|.:||:::.::|.||:
ZIP14       140 QTEEGKPSAIEVWGYGFLCVTVISLCSLMGASVVPFMKKTFYKRLLLYFI  189

ZIP8        167 GLAIGTLFSNAIFQLIPEAFGFNPKIDNYVEKAVAVFGGFYMLFFVERTL  216
                .||||||:|||:||||||||||||: ||||.|:..||||||:.||.|:.|
ZIP14       190 ALAIGTLYSNALFQLIPEAFGFNPQ-DNYVSKSAVVFGGFYLFFFTEKIL  238

ZIP8        217 KMLLKTYGQ--NDHTHFRNDDFGSKEKTHQPKTLPLPAVNGVTCYANPAV  264
                |||||...:  :.|.||.::...||:...:..|..|.  ||...:..|.
ZIP14       239 KMLLKQKNEHHHGHNHFTSETLPSKKDQEEGVTEKLQ--NGDLDHMIPQ-  285

ZIP8        265 TEPNGHIHFD------------TVSVVSLQDGKTEPSSCTCLKGPKLSEI  302
                     |.:.:            .|:.:|:||.:...|:|..|||.:.|:|
ZIP14       286 -----HCNSELDGKAPGTDEKVIVNSMSVQDLQASQSACYWLKGVRYSDI  330

ZIP8        303 GTIAWMITLCDALHNFIDGLAIGASCTLSLLQGLSTSIAILCEEFPHELG  352
                ||:||||||.|.|||||||||||||.|:|:.||:|||:||||||||||||
ZIP14       331 GTLAWMITLSDGLHNFIDGLAIGASFTVSVFQGISTSVAILCEEFPHELG  380

ZIP8        353 DFVILLNAGMSTRQALLFNFLSACSCYVGLAFGILVGNNFAPNIIFALAG  402
                |||||||||||.:|||.|||||||.||:|||||||.|::|:.|.||||||
ZIP14       381 DFVILLNAGMSIQQALFFNFLSACCCYLGLAFGILAGSHFSANWIFALAG  430

ZIP8        403 GMFLYISLADMFPEMNDMLREKVTGRQTDFTF---FMIQNAGMLTGFTAI  449
                ||||||:|||||||||::.:|.    :.:.:|   |:|||.|:||||:.:
ZIP14       431 GMFLYIALADMFPEMNEVCQED----EKNDSFLVPFVIQNLGLLTGFSIM  476

ZIP8        450 LLITLYAGDIELQ    462
                |::|:|:|.|::.
ZIP14       477 LVLTMYSGQIQIG    489
```

—— ZIP8 Predicted transmembrane domains

—— ZIP14 Predicted transmembrane domains

**Supplementary Figure 2. ZIP8 and ZIP14 protein sequence alignment.** Sequence homology analyses showed 46.6% identity and 62.0% similarity between these transporters. Since no crystal structure is available, the predicted transmembrane obtained from UniProtKB sites are indicated as lines in the top (ZIP8) and bottom (ZIP14) of the sequences.

#### Supplementary figure 3

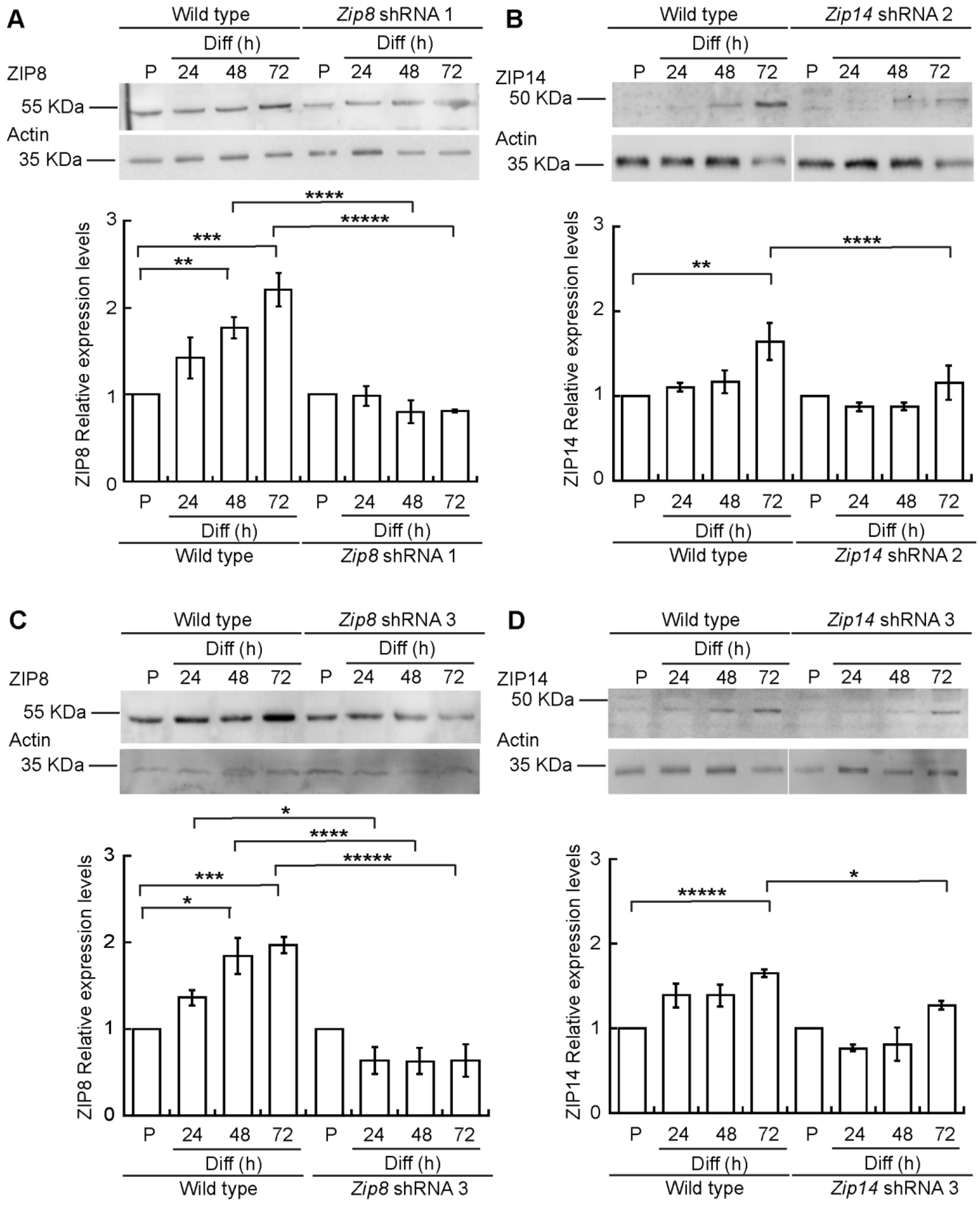


**Supplementary Figure 3. ZIP8 and ZIP14 expression in proliferating and differentiating wild type and additional shRNA knockdown primary myoblasts derived from mouse satellite cells.** ZIP8 and ZIP14 expression in cells transduced with alternate shRNAs utilized to knockdown *Zip8* and *Zip14*. **(A)** Representative immunoblot (top) and quantification (bottom) of ZIP8 levels in proliferating myoblasts and differentiated cells at 24, 48, and 72 h post-differentiation in wild type (left side panel) and *Zip8* shRNA-1-transduced myoblasts (right side panel). **(B)** Representative immunoblot of ZIP14 levels in proliferating and differentiating wild type (left) and *Zip14* shRNA-2-transduced myoblasts. **(C)** Representative immunoblot (top) and quantification (bottom) of ZIP8 levels in proliferating myoblasts and differentiated cells at 24, 48, and 72 h post-differentiation in wild type (left side panel) and *Zip8* shRNA-3-transduced cells (right side panel). **(D)** Representative immunoblot of ZIP14 levels in proliferating and differentiating wild type (left) and *Zip14* shRNA-3-transduced myoblasts. For all samples, shown is mean ± standard error of three independent biological replicates. Immunoblots against actin or Coomasie-stained membranes (Supp. Fig. **8**) were used as loading controls. For wild type differentiating myoblasts, statistical analyses showed significant differences when compared to proliferating cells. Statistical analyses for *Zip8*-knockdown cells showed significant decrease in ZIP8 expression when compared to control cells at the corresponding time points. *Zip14*-knockdown cells showed significant decrease in ZIP14 expression when compared to control cells. *****$P<0.0001$ ****$P<0.001$, ***$P < 0.005$, **$P < 0.01$, *$P \leq 0.05$

**Supplementary figure 4.**

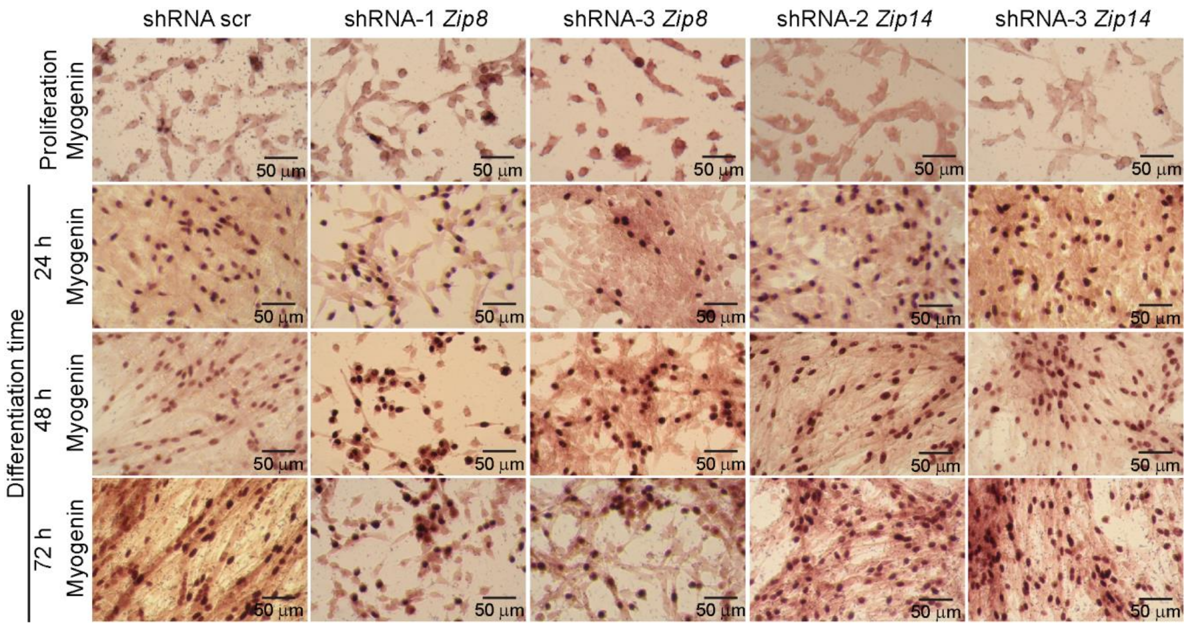


**Supplementary Figure 4. Knockdown of *Zip8* with additional shRNAs impairs differentiation of primary myoblasts.** Representative light micrographs of proliferating and differentiating myoblasts expressing alternate shRNAs against *Zip8* (shRNA-1 and -3 *Zip8*) and *Zip14* (shRNA-2 and -3 *Zip14*) knockdown myoblasts at 24, 48 and 72 h after inducing differentiation. Cells were immunostained with an anti-Myogenin antibody.

**Supplementary figure 5.**

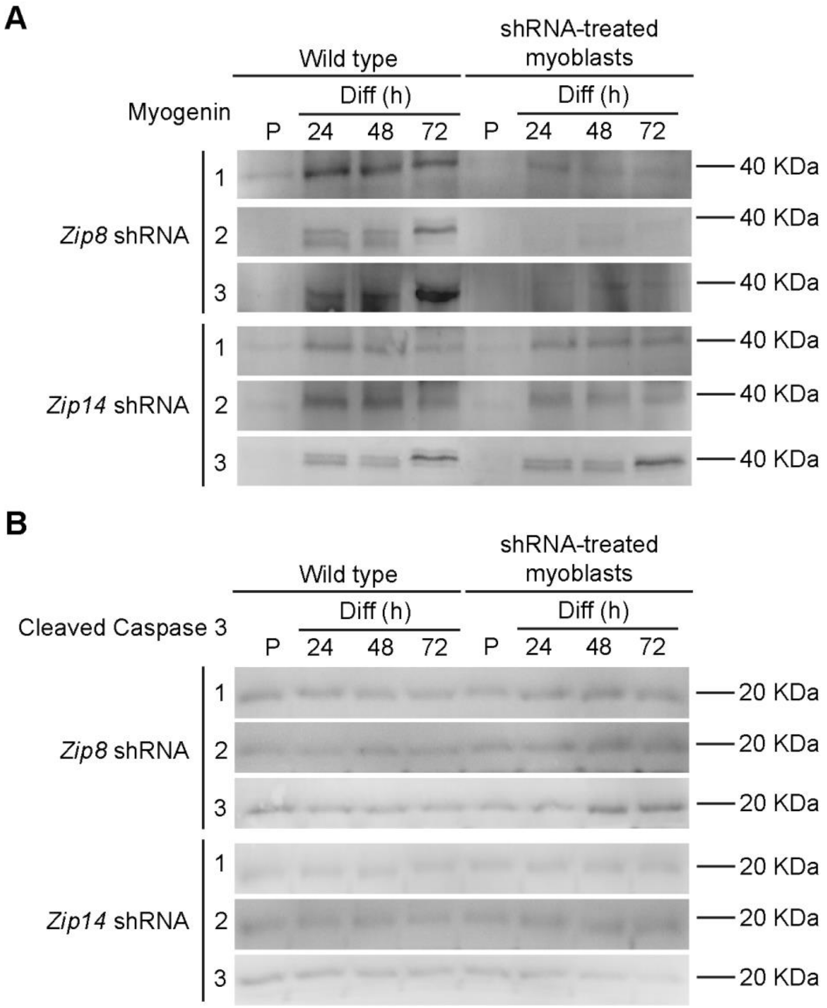


**Supplementary Figure 5. Decreased expression of myogenin and increased activation of caspase 3 is observed in myoblasts exhibiting partial *Zip8* knockdown.** Representative Western blots of the three independent biological replicates of wildtype and each of the three clones of primary myoblasts treated with either with *Zip8* or *Zip14* shRNA developed an anti-myogenin (**A**) and anti-Caspase 3 (**B**) antibodies.

**Supplementary figure 6.**

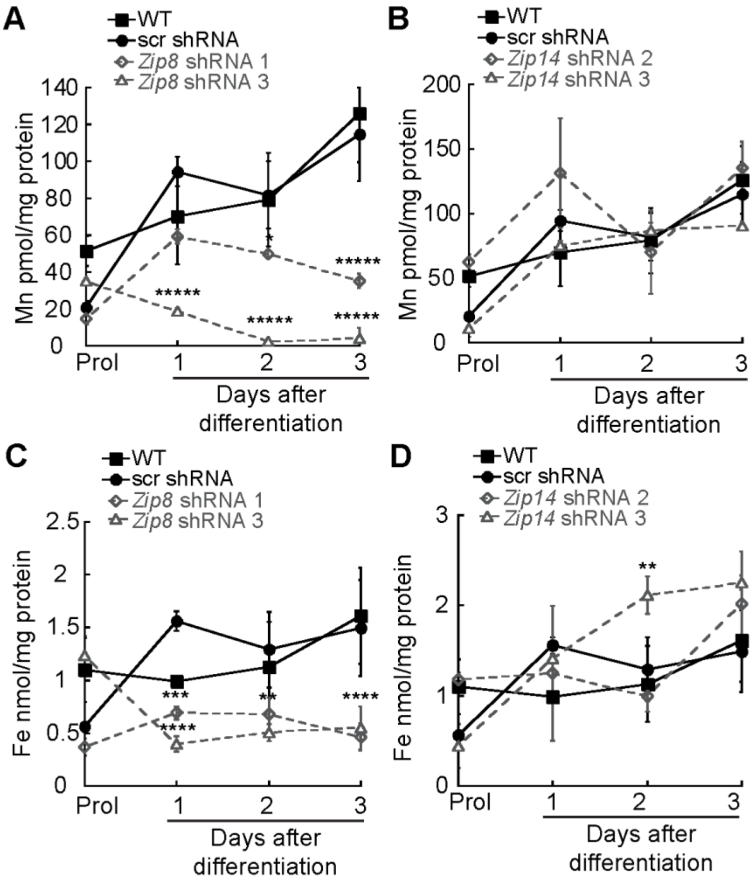


**Supplementary figure 6. Mn and Fe content in *Zip8* and *Zip14* knockdown myoblasts.** ICP-OES analysis of additional metals in wild type, *scr*, *Zip8* and *Zip14* knockdown cells. Whole cell Mn content comparing wild type and scr transduced myoblasts to *Zip8* **(A)** and Zip14 **(B)** partially depleted myoblasts. Whole cell Fe content comparing wild type and scr transduced myoblasts to *Zip8* **(C)** and *Zip14* **(D)** partially depleted myoblasts. Statistical analyses showed significant differences in metal accumulation in differentiating myoblasts when compared to control myoblasts. All data were measured using ICP-OES and normalized to total protein. Shown is mean ± standard error for three biological replicates. ****P<0.001, ***P < 0.005, **P < 0.01, *P ≤ 0.05

**Supplementary Figure 7**

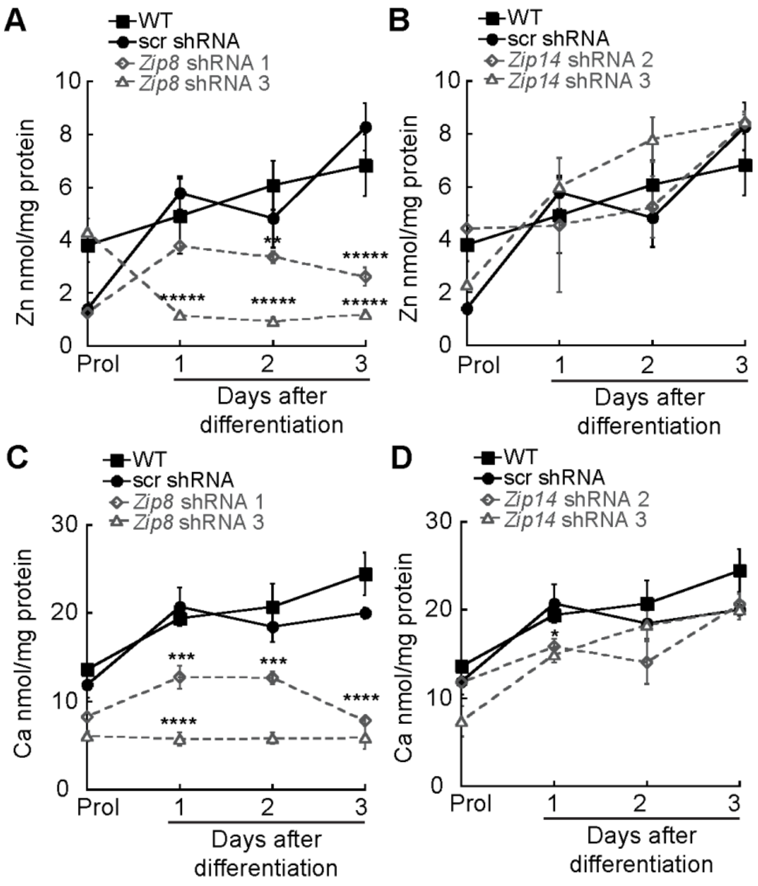


**Supplementary figure 7. Zn and Ca content in *Zip8* and *Zip14* knockdown myoblasts.** ICP-OES analysis of additional metals in wild type, *scr*, *Zip8* and *Zip14* knockdown cells. Whole cell Zn content comparing wild type and scr transduced myoblasts to *Zip8* **(A)** and *Zip14* **(B)** partially depleted myoblasts. Whole cell Ca content comparing wild type and scr transduced myoblasts to *Zip8* **(C)** and *Zip14* **(D)** partially depleted myoblasts. Statistical analyses showed significant differences in metal accumulation in differentiating myoblasts when compared to control myoblasts. All data were measured using ICP-OES and normalized to total protein. Shown is mean ± standard error for three biological replicates. *****P<0.0001 ****P<0.001, ***P < 0.005

### Supplementary Figure 8 (part A)

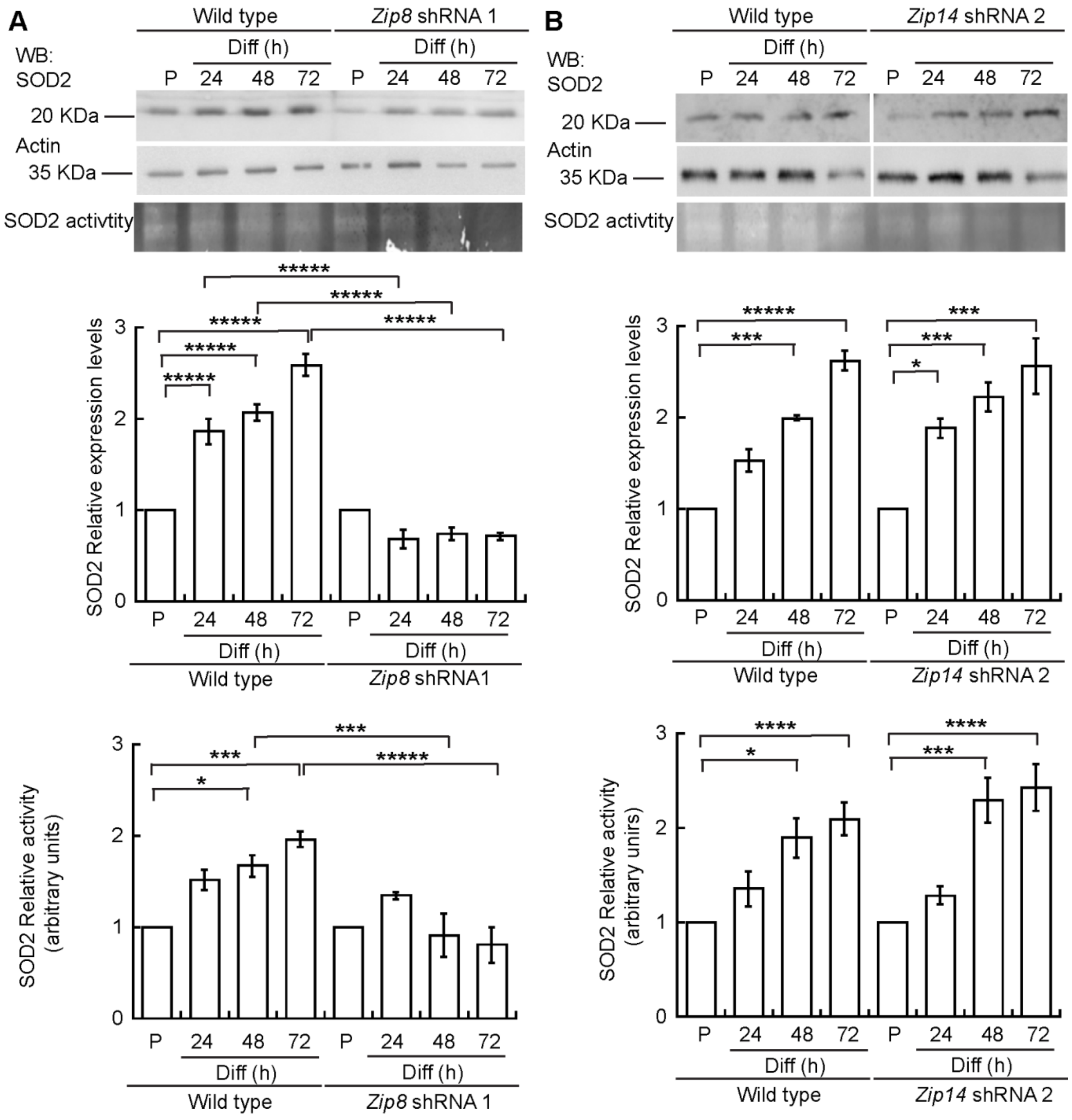

**Supplementary Figure 8 (part B)**

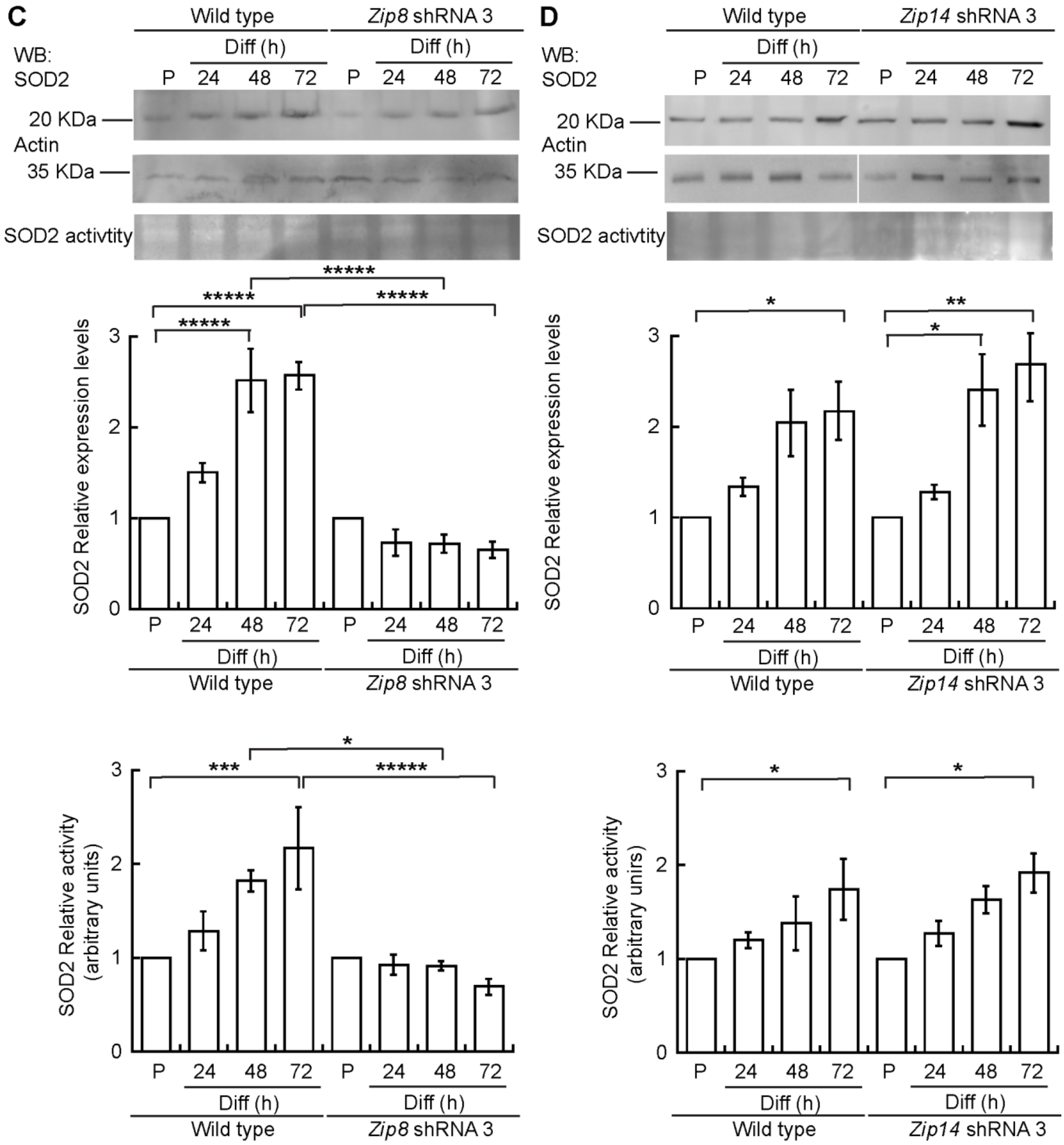


**Supplementary Figure 7. SOD2 expression and activity is decreased in additional *Zip8* but not *Zip14* knockdown primary myoblasts.** Additional immunoblots and activity gels analyses for SOD2 in proliferating and differentiating myoblasts expressing alternate shRNAs. **(A)** Representative Western blots and activity gels (top panels) and quantification of SOD levels and activity (bottom panels) in wild type and cells expressing an alternate *Zip8* shRNA (shRNA-1 *Zip8*). **(B)** Representative Western blots and activity gels (top panels) and quantification of SOD levels and activity (bottom panels) in wild type and cells expressing an alternate *Zip14* shRNA (shRNA-2 *Zip14*). **(C)** Representative Western blots and activity gels (top panels) and quantification of SOD levels and activity (bottom panels) in wild type and cells expressing an alternate *Zip8* shRNA (shRNA-3 *Zip8*). **(D)** Representative Western blots and activity gels (top panels) and quantification of SOD levels and activity (bottom panels) in wild type and cells expressing an alternate *Zip14* shRNA (shRNA-3 *Zip14*). For all samples, blots against actin and Coomasie-stained membranes were used as loading controls (Supp. Figure **9**). Shown is mean ± standard error for three biological replicates. For wild type differentiating myoblasts, statistical analyses showed significant differences when compared to proliferating cells. Statistical analyses for *Zip8*-knockdown cells showed significant differences when compared to control cells at the corresponding time points. Statistical analyses for *Zip14*-knockdown cells showed significant differences in SOD2 levels and activity when compared to proliferating *Zip14*-mutant myoblasts. *****P<0.0001 ****P<0.001, ***P < 0.005, **P < 0.01, *P ≤ 0.05

**Supplementary Figure 9.**

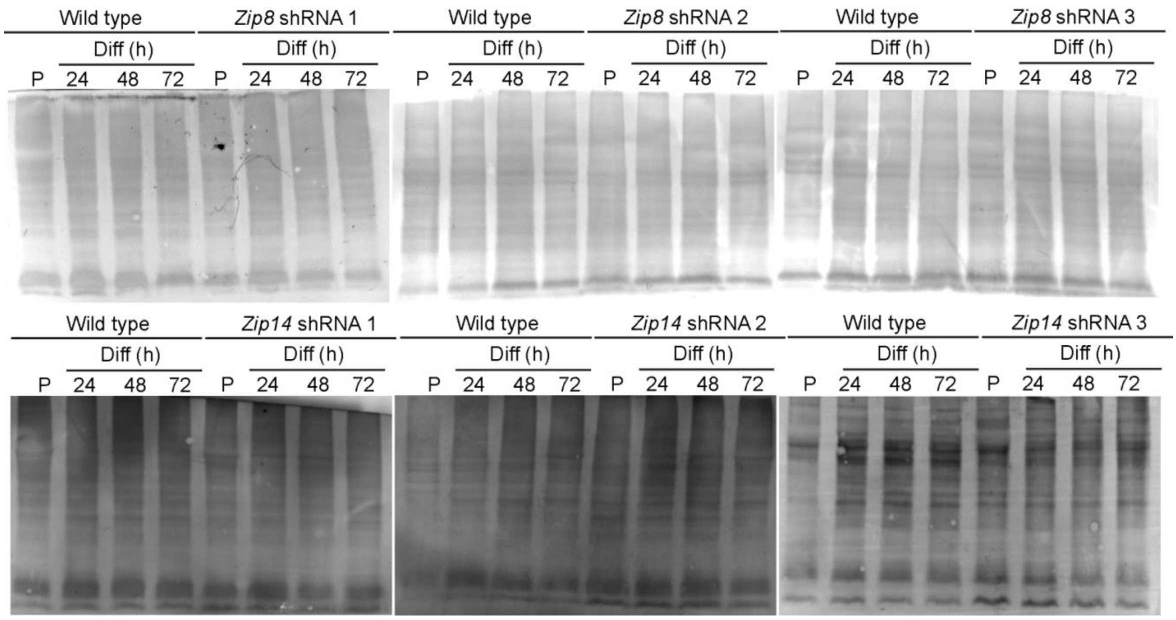


**Supplementary Figure 9. Coomassie-stained immunoblot membrane controls.** Representative Coomasie-stained membranes demonstrating total protein load. These membranes were utilized as a loading control for in-gel SOD2 activity assays shown in Fig. **6** and Supp. Fig. **6**.

`

**Supplementary figure 10**

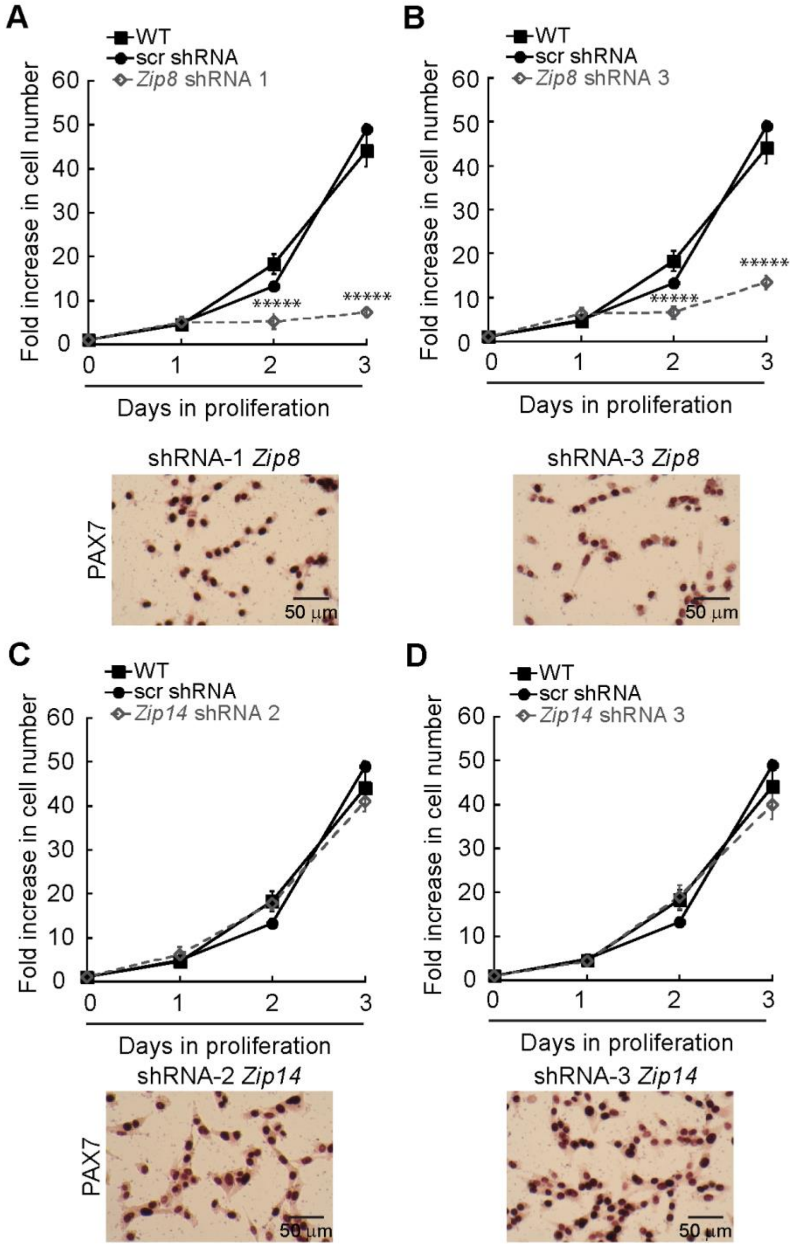


**Supplementary figure 10. Partial depletion of *Zip8*, but not *Zip14*, impairs growth of primary myoblasts.** Cell counting assay of proliferating wild type myoblasts, and cells transduced with scrambled shRNA (shRNA scr), *Zip8* (**A, B**) or *Zip14* shRNAs (**C, D**). Data in A-C are mean ± standard error for three independent experiments. *****P<0.0001. Representative light micrographs of *Zip8* (**A-B**, lower panel) and *Zip14* (**C-D** lower panel) knockdown proliferating myoblasts immunostained with an anti-Pax7 antibody.

**Supplementary figure 11.**

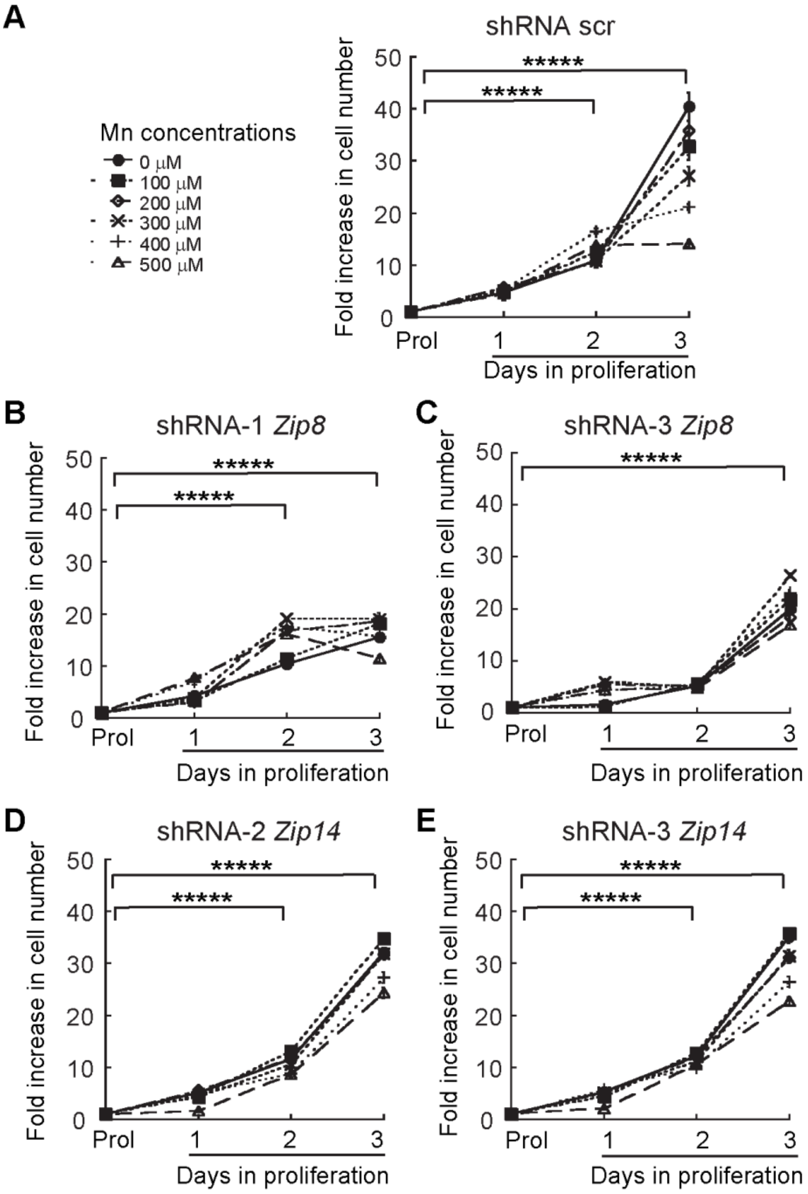


**Supplementary Figure 4. Mn does not rescue the growth defect observed in additional knockdowns of *Zip8* primary myoblasts.** Cell counting assays performed in myoblasts expressing alternate shRNAs grown in medium supplemented with exogenous Mn. **(A)** Cell counting assay of proliferating myoblasts expressing scrambled shRNAs (shRNA scr) grown in increasing concentrations of Mn. Statistical analyses showed significant differences when compared to the proliferating cells. Cell counting assay of shRNA-1 (**B**), and shRNA-3 (**C**) *Zip8* knockdown myoblasts grown in increasing concentrations of Mn. Statistical analyses showed significant differences when compared to control cells at the corresponding proliferation or differentiation time points. Cell counting assay of shRNA-2 (**D**), and sRNA-3 (**E**) of *Zip14* knockdown myoblasts grown in increasing concentrations of Mn. Statistical analyses showed significant differences when comparing differentiating myoblasts to proliferating cells. For all experiments, data represent mean ± standard error for three biological replicates. *****P<0.0001

**SUPPLEMENTARY TABLE**

**Supplementary table 1. Sequences of shRNA used.**

| **Name** | **Sequence** |
|---|---|
| Zip8 sh1 | CCGGGCCAAGTTATCTCAGGAATTACTCGAGTAATTCCTGAGATAACTTGGCTTTTTG |
| Zip8 sh2 | CCGGCAACGCGGGAAGGCATTTAATCTCGAGATTAAATGCCTTCCCGCGTTGTTTTTG |
| Zip8 sh3 | CCGGTACGCAGGAGACATCGAATTGCTCGAGCAATTCGATGTCTCCTGCGTATTTTTG |
| Zip14 sh1 | CCGGCCTCCCTTTCTCTTGGAAGAACTCGAGTTCTTCCAAGAGAAAGGGAGGTTTTTG |
| Zip14 sh2 | CCGGCCTCTACTCCAACGCCCTCTTCTCGAGAAGAGGGCGTTGGAGTAGAGGTTTTTG |
| Zip14 sh3 | CCGGTCCAGAATCTTGGCCTCCTAACTCGAGTTAGGAGGCCAAGATTCTGGATTTTTG |

#### SUPPLEMENTARY METHODS

##### Alignments

Sequences for ZIP8 and ZIP14 were obtained from UniProt and were aligned using EMBOSS Needle Pairwise Sequence Alignment software [1,2].

##### Membrane staining by Coomassie Brilliant Blue.

After developing the PVDF membranes with the appropriate antibodies, the membranes were rinse with $H_2O$ for 10 min. Then, they were stain with 0.025% (w/v) Coomassie brilliant blue R-250 (Sigma) in 40% methanol/7% acetic acid (v/v) for 5 min. Membranes were de-stained by washing 3 times with 50% methanol/7% acetic acid for 10 min and rinsed with $H_2O$ [3].
